# Strategies for vaccine design for corona virus using Immunoinformatics techniques

**DOI:** 10.1101/2020.02.27.967422

**Authors:** Anamika Basu, Anasua Sarkar, Ujjwal Maulik

**Affiliations:** Assistant Professor, Gurudas College, India; Computer Science and Engineering Department, Jadavpur University

**Keywords:** Immunoinformatics, vaccine design, Coronavirus, nonstructural protein 4 of beta coronavirus, B cell epitope, T cell epitope, molecular docking

## Abstract

The cutting-edge technology vaccinomics is the combination of two topics immunogenetics and immunogenomics with the knowledge of systems biology and immune profiling for designing vaccine against infectious disease. In our present study, an epitope-based peptide vaccine against nonstructural protein 4 of beta coronavirus, using a combination of B cell and T cell epitope predictions, followed by molecular docking methods are performed. Here, protein sequences of homologous nonstructural protein 4 of beta coronavirus are collected and conserved regions present in them are investigated via phylogenetic study to determine the most immunogenic part of protein. From the identified region of the target protein, the peptide sequence IRNTTNPSAR from the region ranging from 38-47 and the sequence PTDTYTSVYLGKFRG from the positions of 76-90 are considered as the most potential B cell and T cell epitopes respectively. Furthermore, this predicted T cell epitopes PTDTYTSVY and PTDTYTSVYLGKFRG interacted with MHC allelic proteins HLA-A*01:01 and HLA-DRB5*01:01 respectively with the low IC_50_ values. These epitopes are perfectly fitted into the epitope binding grooves of alpha helix of MHC I molecule and MHC II molecule with binding energy scores −725.0 Kcal/mole and −786.0 Kcal/mole respectively, showing stability in MHC molecules binding. This MHC restricted epitope PTDTYTSVY also showed a good conservancy of 50.16% in world population coverage. This MHC I HLA-A*01:01 allele is present among 58.87% of Chinese population also. Therefore, the epitopes IRNTTNPSAR and PTDTYTSVYLGKFRG may be considered as potential peptides for peptide-based vaccine for coronavirus after further experimental study.

## 1. Introduction

According to World Health Organization, Coronaviruses (CoV) are a large family of RNA viruses that cause infection ranging from the common cold to more severe diseases such as Middle East Respiratory Syndrome (MERS-CoV) and Severe Acute Respiratory Syndrome (SARS-CoV). A novel coronavirus (nCoV) is a new strain that has not been previously identified in humans. 2019 Novel Coronavirus (2019-nCoV) is a coronavirus identified as the cause of an outbreak of respiratory illness first detected in Wuhan, China.

Coronaviruses are zoonotic, meaning they are transmitted between animals and people. Detailed investigations found that SARS-CoV has been transmitted from civet cats to humans and MERS-CoV from camels to humans. Several known coronaviruses are circulating in animals that have not yet infected humans. Early on, many of the patients in the outbreak in Wuhan, China reportedly have some link to a large seafood and animal market, suggesting animal-to-person spread. However, a growing number of patients reportedly have not had exposure to animal markets, indicating person-to-person spread is occurring. The 2019-nCoV is spreading from person to person in China and limited spread among close contacts has been detected in some countries outside China. Common signs of infection include respiratory symptoms, fever, cough, shortness of breath and breathing difficulties. In more severe cases, infection can cause pneumonia, severe acute respiratory syndrome, kidney failure and even death. There is currently no vaccine to protect against 2019-nCoV.

According to Lu et al, 2020 [1], genome sequence of 2019-nCoV is closely related (with 88% identity) to two bat-derived severe acute respiratory syndrome (SARS)-like coronaviruses, bat-SL-CoVZC45 and bat-SL-CoVZXC21 and genetically distinct from SARS-CoV. Two complete virus genomes (HKU-SZ-002a and HKU-SZ-005b) are sequenced from 2019-nCoV infected patients [2]. HKU-SZ-002a and HKU-SZ-005b differ from each other by only two bases. One of them is a non-synonymous mutation at amino acid position 336 of non-structural protein 4 (Ser336 for HKU-SZ-002a; Leu336 for HKU-SZ-005b). The amino acid sequence of the N-terminal domain of Spike subunit 1 of this novel coronavirus is around 66% identical to those of the SARS-related coronaviruses, and the core domain of the receptor binding domain of this novel coronavirus has about 68% amino acid identity with those of the SARS-related coronavirus. But the protein sequence of the external subdomain region of receptor binding domain of Spike subunit 1 has only 39% identity, which might affect the choice of human receptor and therefore the biological activity of this virus.

Coronaviruses encode large replicase polyproteins which are proteolytically processed by viral proteases to generate mature nonstructural proteins (nsps) that form the viral replication complex. Positive-strand RNA viruses, such as coronaviruses, can induce cellular membrane rearrangements during replication to form replication organelles which allows efficient viral RNA synthesis. Nonstructural Protein 4 alone induces membrane pairing in infectious bronchitis virus [3]. Infection with coronavirus causes rearrangement in the host cell membrane to accumulate a replication and transcription complex in which replication of the viral genome and transcription of viral mRNA can occur. For coronaviruses, a major pathogenicity factor has now been identified with non-structural protein 1 in a murine model of coronavirus infection [4]. Sakai et al in 2017 [5] highlighted the role of nsp4 of SARS coronavirus in viral replication. It has been discovered that only a change in two amino acids in nsp4 protein sequence can abolish viral replication completely. So, to prevent coronavirus infection by designing vaccine, nonstructural protein 4 of beta coronavirus can be selected as target protein in this study.

Prevention of viral diseases by vaccination purposes for controlled induction of protective immune responses against viral pathogens such as small pox, hepatitis etc. Attenuated viral vaccines can be produced by targeting essential pathogenicity factors. This study has implicated for the rational design of live attenuated coronavirus vaccines aiming to prevent coronavirus-induced diseases of human as well as animal origin, including the lethal severe acute respiratory syndrome.

In this study nonstructural protein 4 has been analyzed in order to predict informative epitopes which will help for future vaccine designing. The knowledge of peptide vaccine is used to identify B-cell and T-cell epitopes which can induce specific immune responses [6], [7]. These designed epitopes will offer cost effective, high quality therapeutics against corona virus. By using online algorithms in immunoinformatics, potentially active immunogenic T and B cell epitopes for this viral protein has been identified and modelled to design possible peptide vaccine for coronavirus infection.

## 2. Materials and methods

### 2.1 Retrieval of Non-structural protein NS4 in Beta coronavirus HKU24

The protein family information of Coronavirus nonstructural protein NS4 containing 78 protein is retrieved from InterPro database (https://www.ebi.ac.uk/interpro/entry/InterPro/IPR005603/). To find conserved region, retrieved sequences were aligned using Muscle tool 3.8.31 [8] where *k*-mer clustering is used. In the next step a phylogenetic tree is constructed by the method known as progressive alignment. The evolutionary divergence analysis for all 78 virus proteins are completed by forming a phylogenetic tree using Phy ML 3.1/3.0 aLRT software [9]. Here the phylogenetic tree was reconstructed using the maximum likelihood method. The default substitution model was selected assuming an estimated proportional of invariant sites (of 0.008) and 4 gamma-distributed rate categorized to account for rate heterogenicity across sites. The gamma shape parameter was estimated directly from the data (gamma = 1.788). Reliability for internal branch was assessed using the aLRT test (SH-like).

Among these proteins non-structural protein NS4, with accession number A0A0A7UXD8, known as Beta coronavirus HKU24 (due to its Chinese origin) with length 136 is selected in FASTA format.

### 2.2 Protein antigenicity determination

Antigenicity of this protein is predicted by VaxiJen v 2.0, an online prediction server [10].

### 2.3 Potential B cell epitope prediction

B cell epitopes are part of allergen which come in interaction with B lymphocytes to induce immune response. Among two types of B cell epitopes, linear type of B cell epitopes is predicted.

Various physico-chemical properties e.g. hydrophilicity, flexibility, accessibility, turns, exposed surface, polarity and antigenic propensity of peptides chains have been estimated to identify the locations of linear epitopes of an antigenic protein [11]. Thus, different tools from IEDB (www.iedb.org), including the classical propensity scale methods such as Kolaskar and Tongaonkar antigenicity scale [12], E mini surface accessibility prediction [13], Parker hydrophilicity prediction [14], Karplus and Schulz flexibilty prediction [15], Bepipred linear epitope prediction [16] and Chou and Fashman beta turn prediction tool [17] are used to predict linear or continous B cell epitopes of nonstructural protein NS4 protein in coronavirus. With the help of graphical findings and prediction scores the most probable B cell epitope of that antigenic protein has been identified. BepiPred prediction method is a combination of Hidden Markov model and propensity scale method to predict score and identification of B cell epitopes of antigenic protein [16].

### 2.4 Potential T-cell epitope prediction

#### 2.4.1 MHC I T cell epitope prediction

Linear T-cell epitopes for MHC-I binding for nonstructural protein NS4 protein in coronavirus are recognized by consensus methods using various methods such as Artificial neural network (ANN) [18], Stabilized matrix method (SMM) [19] and Scoring Matrices Derived from Combinatorial Peptide Libraries (Comblib) from tools for MHC -I binding prediction methods of Immune Epitope Database (IEDB) (www.iedb.org) [20]. This server-based method forecasts the MHC class I binding predication to 26 MHC supertypes as percentile rank. SMM algorithm of MHC-1 binding, transporter of antigenic peptides (TAP) transport efficiency and proteasomal cleavage efficiency are also considered to determine the IC_50_ values for processing prediction of epitopes by MHC-I molecules [21]. On the basis of low IC_50_ values, 5 best epitopes bind with specific MHC-1 molecules are elected for further evaluation.

#### 2.4.2 MHC II T cell epitope prediction

CD4+ T-cell receptor responses against concerned nonstructural protein NS4 protein are done by using Peptide binding to MHC class II molecules software using MHC II binding prediction tool in IEDB analysis resource, including a consensus approach which combines NN-align, SMM-align [21] and Combinatorial library methods. For this prediction we chose thirty HLA class II alleles from the reference set. For the predicted T-cell epitopes with low percentile rank are identified and their IC_50_ values for respective alleles are determined by SMM-align method [22].

### 2.5 Analysis of population coverage

Population coverage for identified T cell epitopes is assessed for world, as well as China and Indian population with the help of IEDB population coverage calculation tool [23]. This tool calculates the fraction of individuals predicted to reply to a specified set of epitopes with recognized MHC restrictions. This calculation is finished considering the HLA genotypic frequencies assuming non-linkage disequilibrium between HLA loci.

### 2.6 Docking study of T cell epitopes

For docking studies, the T cell epitope PTDTYTSVY (MHC I restricted) and PTDTYTSVYLGKFRG (MHC II restricted) are selected and subjected to PEP-FOLD server [24], [25] for 3D structure formation. To identify the molecular interactions with specific HLA protein for respective epitopes, docking studies are performed with ClusPro 2.2 web server [26]. Cluster scores for lowest binding energy prediction are calculated using the formula-E = 0.40E_{rep} + −0.40E_{att} + 600E_{elec} + 1.00E_{DARS}. Here, repulsive, attractive, electrostatic as well as interactions extracted from the decoys as the reference state, are considered for structure-based pairwise potential calculation in epitope-MHC molecule docking [27]. Modified PDB ID 4NQX for HLA-A*01:01 is used as allelic protein for docking study of T cell epitope with MHC I restricted. Similarly, 3D structure (PDB ID 1FV1) for HLA-DRB5*01:01 MHC molecule is used as receptor molecule for T cell epitope docking analysis with MHC II restriction.

## 3 Results

### 3.1 Retrieval of Non-structural protein NS4 in Beta coronavirus HKU24

From InterPro database, 78 homologous protein sequences are recovered for nonstructural protein NS4 protein for coronavirus as shown in Table 1.

**Table 1.**
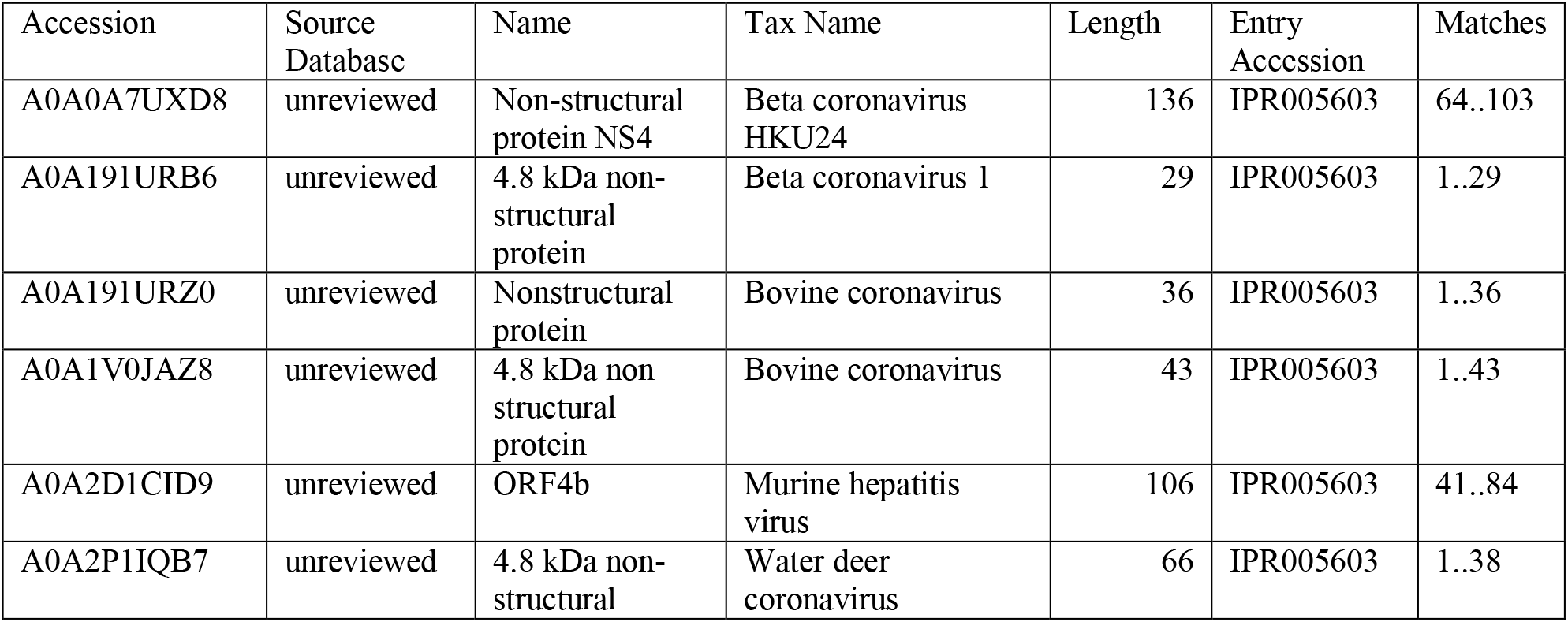
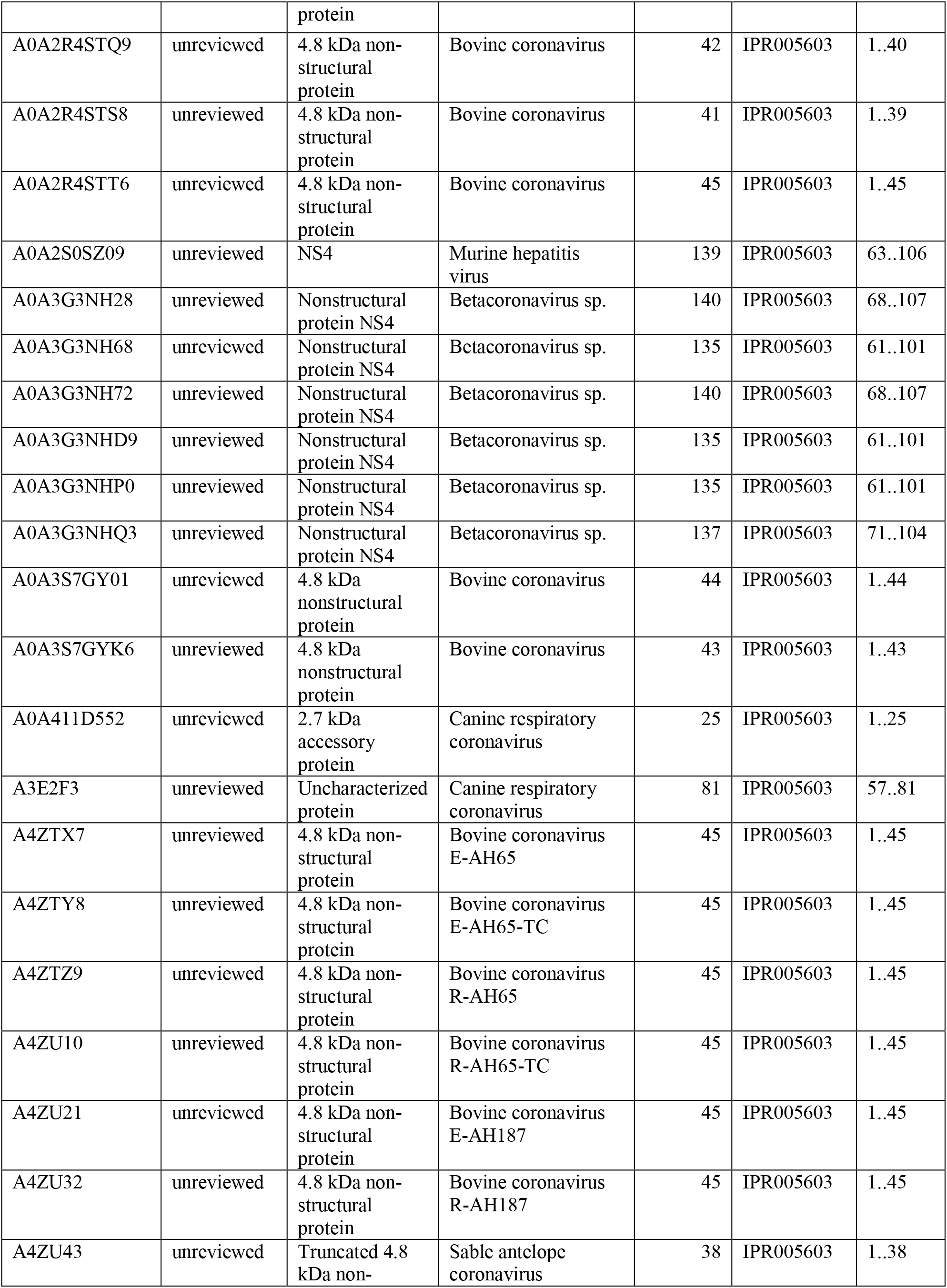
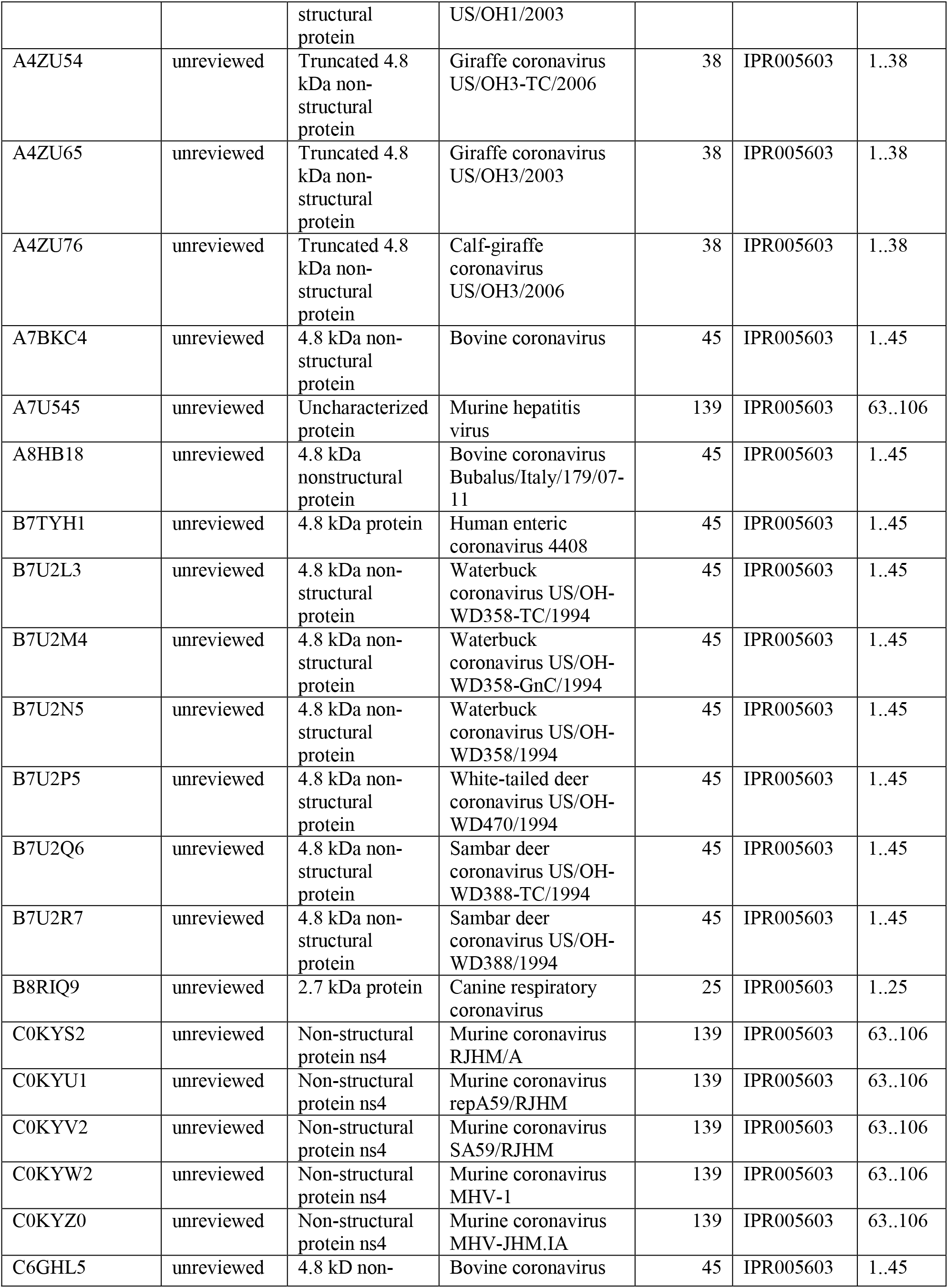
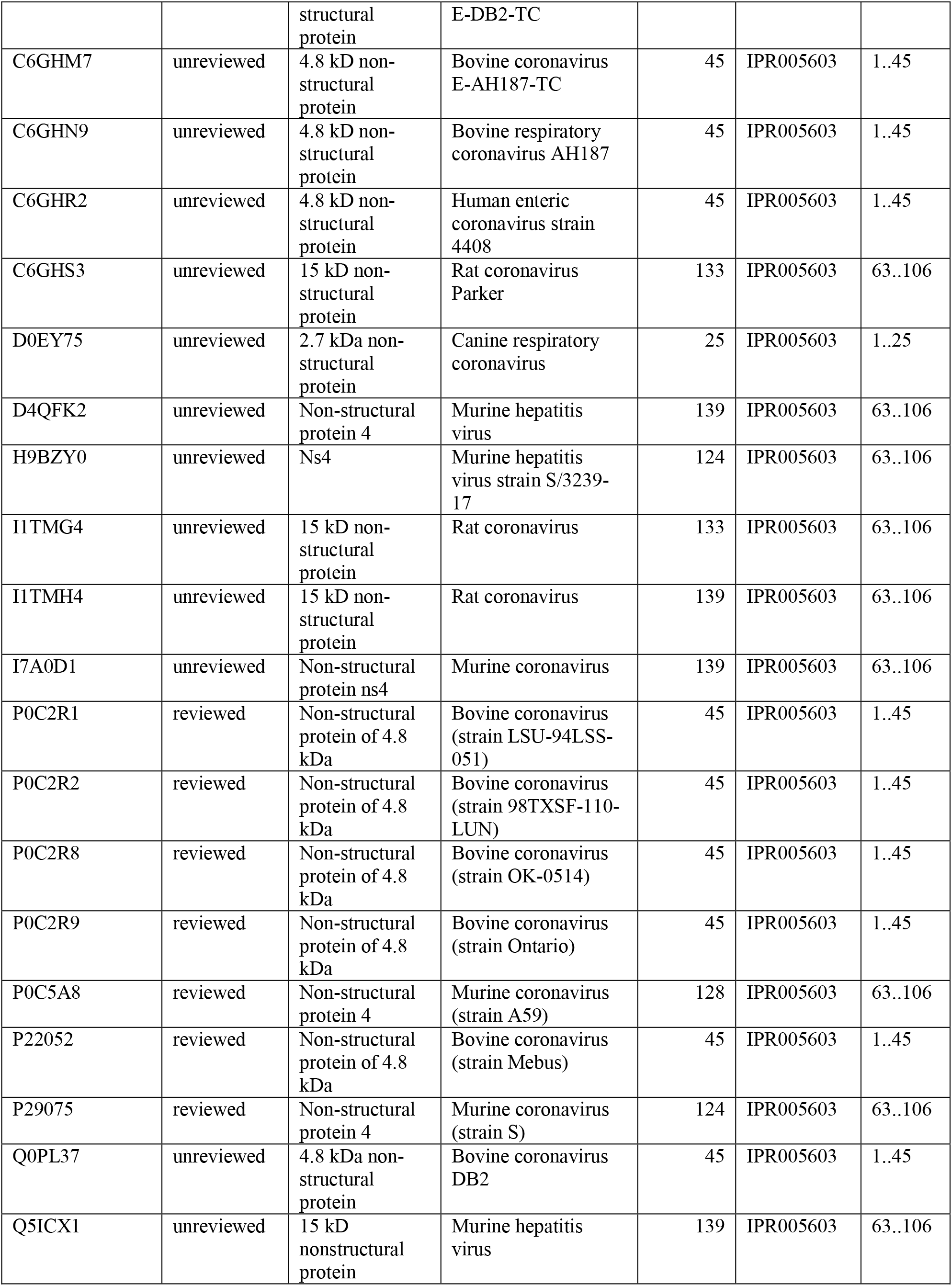
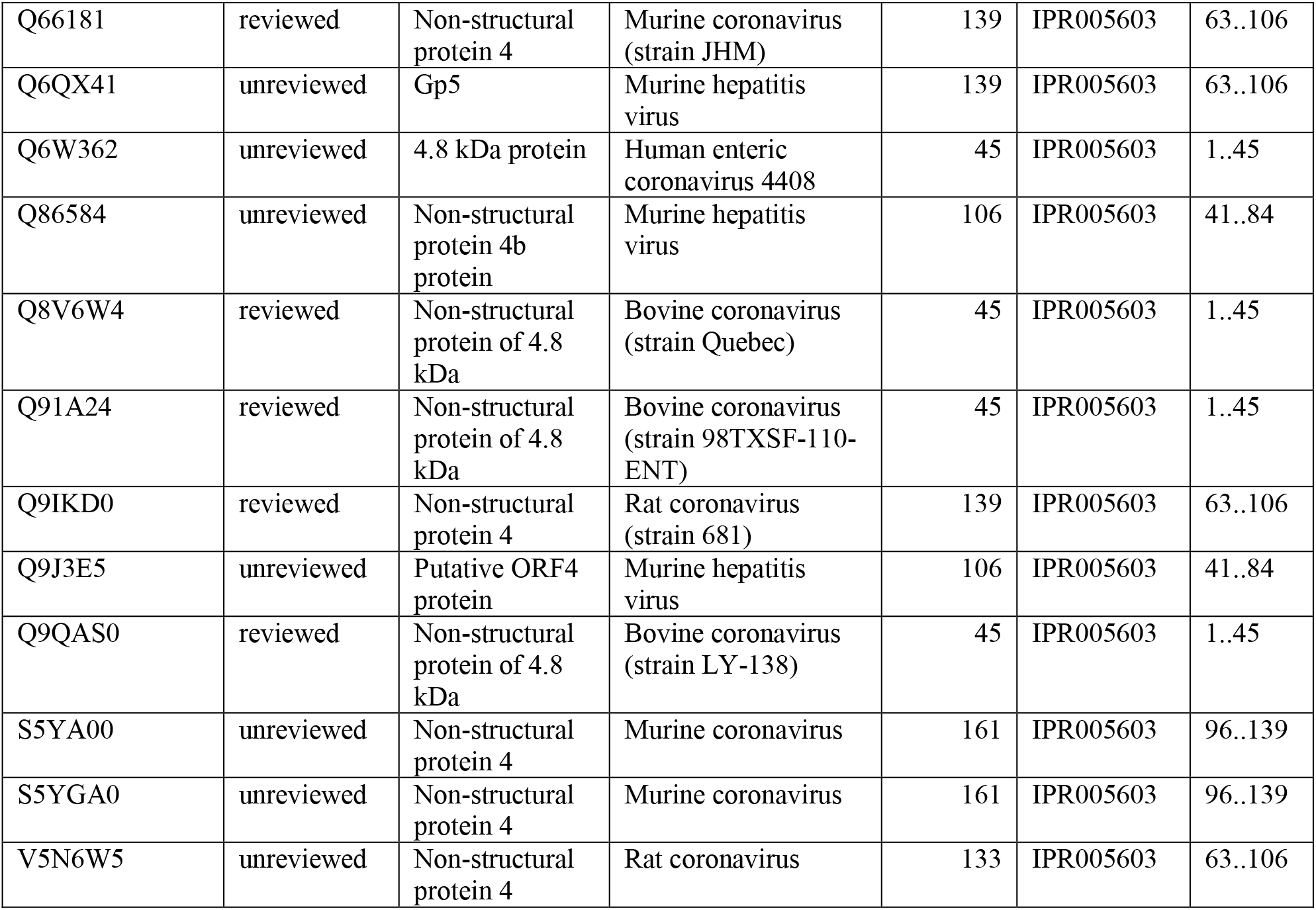
List of 78 nonstructural protein NS4 protein for coronavirus

A phylogenetic tree illustrating the evolutionary relationship among the 78 homologous non-structural protein 4 of coronavirus is depicted in Figure 1a and the relevant enlarged version is shown in Figure 1b.

**Figure 1a.**
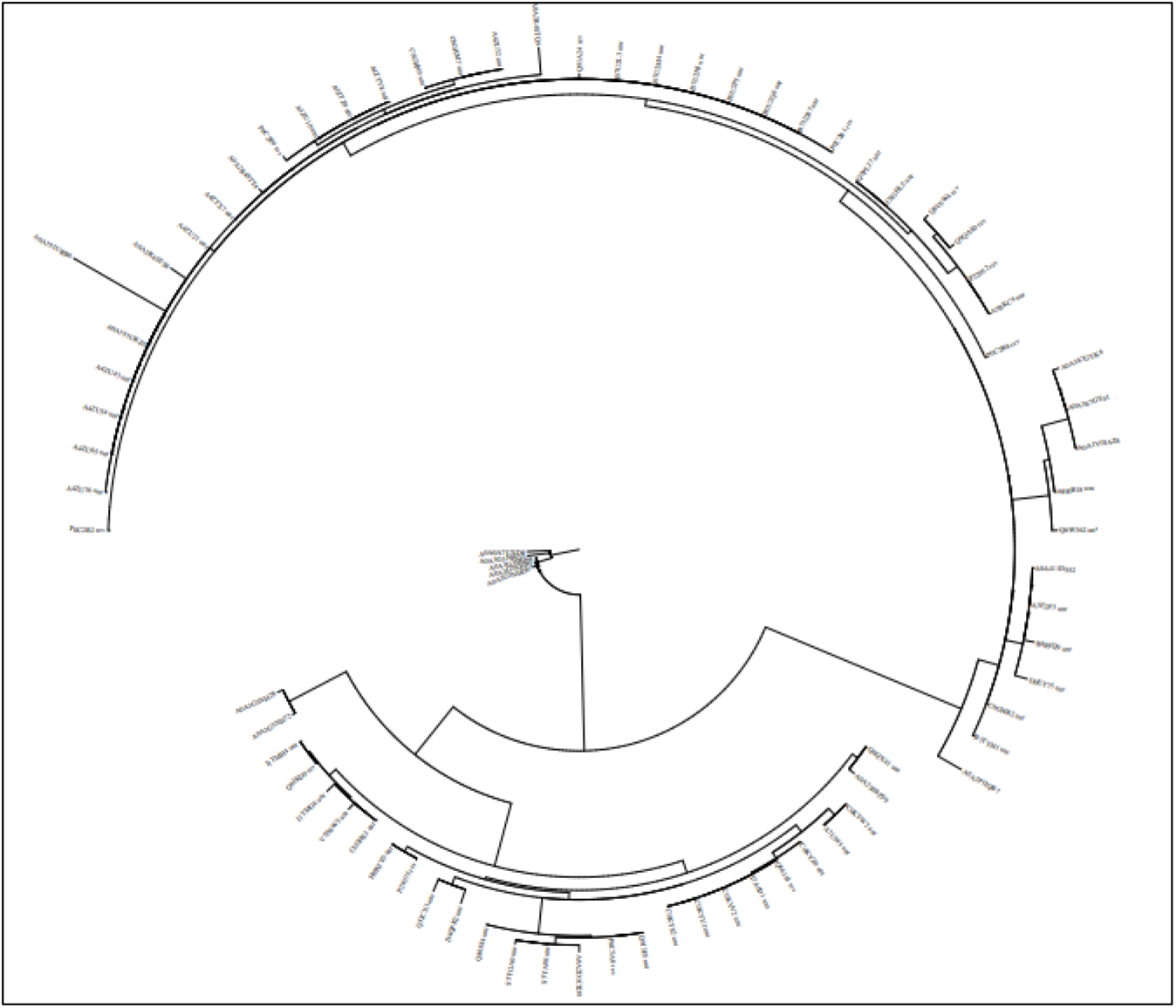
A phylogenetic tree of non-structural protein NS4

**Figure 1b.**
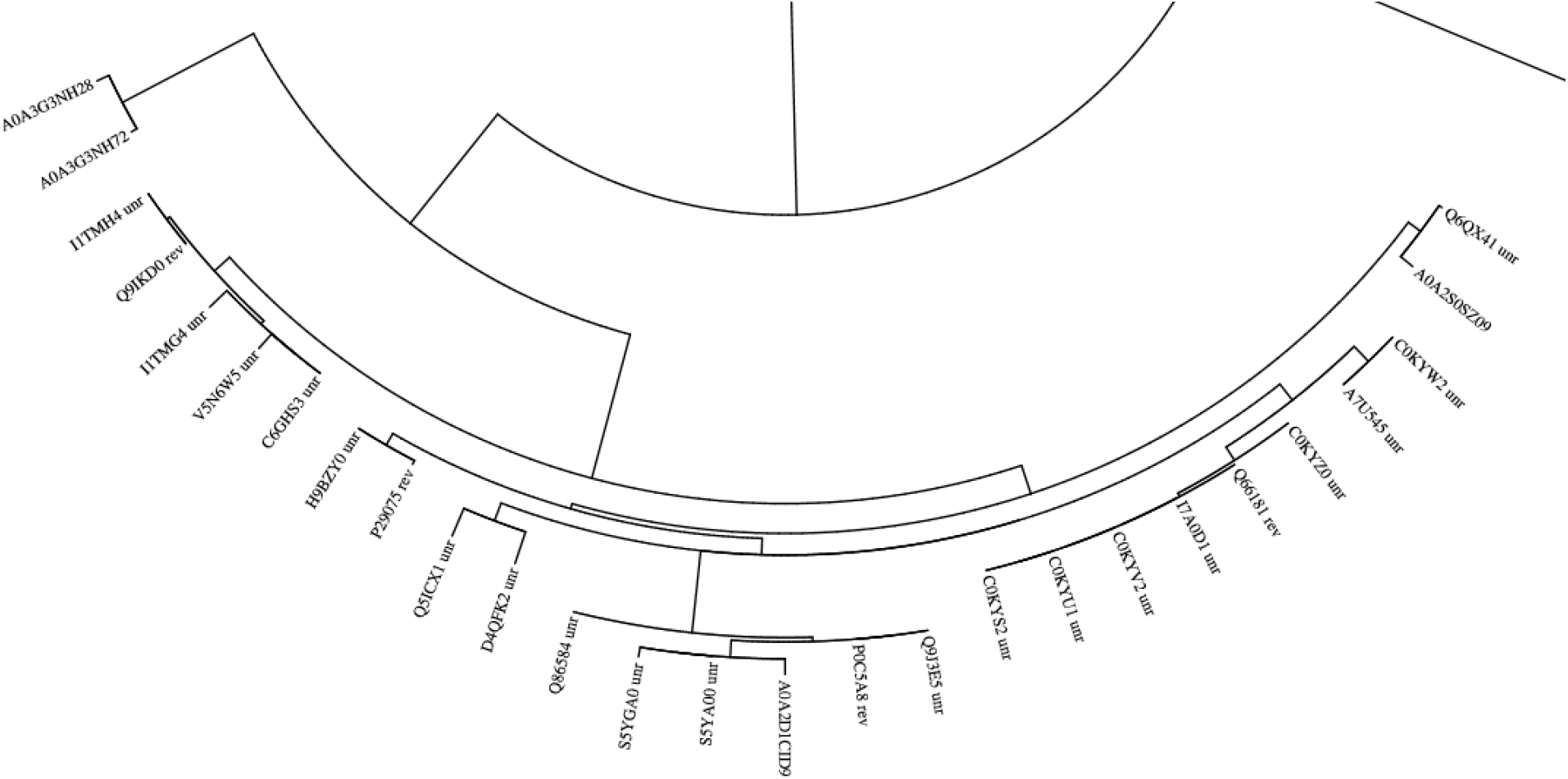
An enlarged version of phylogenetic tree of non-structural protein NS4

Phylogenetic tree shows that Non-structural protein NS4 of Beta coronavirus HKU24 (AOAO7UXD8_9BETC), rat coronavirus (NS4_CVRSD), murine coronavirus (NS4_CVMJH, NS$_CVMS and NSA_CVMA5) and ORF4 protein of murine hepatitis virus (Q9J3EC) are very similar in their structures (identity vary 42.3% to 38.3%). All these five protein sequences are aligned to identify conserved sequences with varying length using Muscle tool 3.8.31[8] is shown in Figure 2. Conserved regions are highlighted.

**Figure 2.**
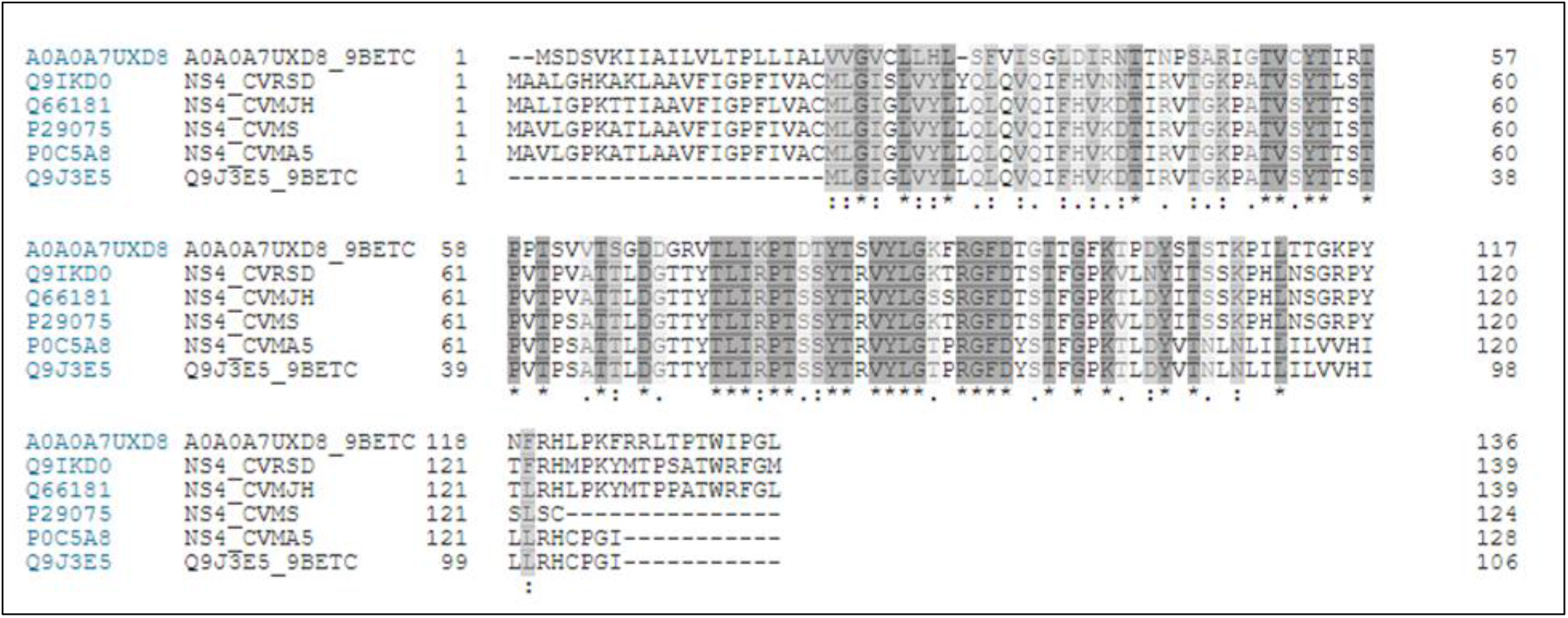
Multiple sequence alignment five Non-structural protein NS4

### 3.2 Potential T-cell epitope prediction

#### 3.2.1 MHC I T cell epitope prediction

#### 3.2.2 MHC II T cell epitope prediction

Among the five peptide sequences, ^76^ PTDTYTSVY ^84^ peptide sequence (MHC I restricted) and ^76^ PTDTYTSVYLGKFRG ^90^ (MHC II restricted) are selected as most probable T cell epitope for the antigenic protein present in non-structural protein NS4 of beta coronavirus HKU24 on the basis of its interactions with highest number of alleles (Table 2 and 3). The IC_50_ values of these peptides are least for the MHCI HLA-A*01:01and MHC II allele HLA-DRB5*01:01 respectively.

**Table 2.**
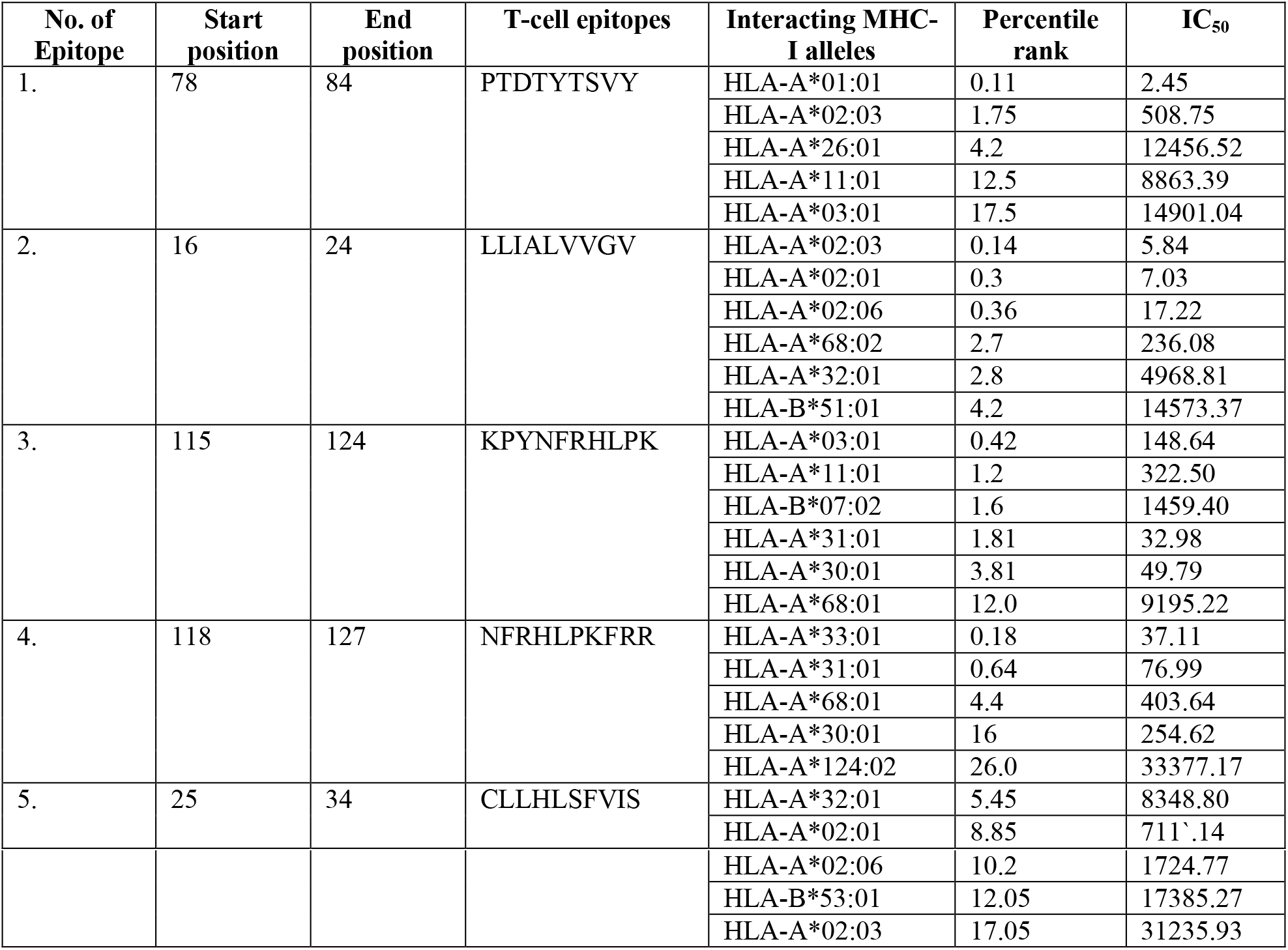
CD8+ T cell epitopes

**Table 3.** CD4+ t cell epitopes

| No. of Epitope | Start position | End position | T-cell epitopes | Interacting MHC-II alleles | Percentile rank |
| --- | --- | --- | --- | --- | --- |
| 1. | 115 | 129 | KPYNFRHLPKFRRLT | HLA-DRB5*01:01 | 0.04 |
|  |  |  |  | HLA-DRB1*15:01 | 16.00 |
|  |  |  |  | HLA-DRB3*02:02 | 20.00 |
|  |  |  |  | HLA-DRB3*01:01 | 31.00 |
|  |  |  |  | HLA-DRB1*03:01 | 34.00 |
|  |  |  |  | HLA-DRB4*01:01 | 40.00 |
|  |  |  |  | HLA-DRB1*07:01 | 42.00 |
| 2. | 16 | 30 | LLIALVVGVCLLHLS | HLA-DRB1*15:01 | 6.00 |
|  |  |  |  | HLA-DRB1*07:01 | 7.70 |
|  |  |  |  | HLA-DRB1*03:01 | 27.00 |
|  |  |  |  | HLA-DRB 4*01:01 | 27.00 |
|  |  |  |  | HLA-DRB 5*01:01 | 37.00 |
|  |  |  |  | HLA-DRB 3*01:01 | 62.00 |
|  |  |  |  | HLA-DRB 3*02:02 | 78.00 |
| 3. | 118 | 132 | NFRHLPKFRRLTPTW | HLA-DRB 5*01:01 | 0.25 |
|  |  |  |  | HLA-DRB 1*15:01 | 11.00 |
|  |  |  |  | HLA-DRB 3*02:02 | 29.00 |
|  |  |  |  | HLA-DRB 4*01:01 | 29.00 |
|  |  |  |  | HLA-DRB 1*03:01 | 52.00 |
|  |  |  |  | HLA-DRB 3*01:01 | 70.00 |
|  |  |  |  | HLA-DRB 1*07:01 | 71.00 |
| 4. | 76 | 90 | PTDTYTSVYL GKFRG | HLA-DRB 5*01:01 | 22.00 |
|  |  |  |  | HLA-DRB 1*15:01 | 24.00 |
|  |  |  |  | HLA-DRB 3*01:01 | 35.00 |
|  |  |  |  | HLA-DRB 1*07:01 | 41.00 |
|  |  |  |  | HLA-DRB 3*02:02 | 69.00 |
| 5. | 25 | 39 | CLLHLSFVISGLDIR | HLA-DRB 1*07:01 | 6.10 |
|  |  |  |  | HLA-DRB 1*03:01 | 7.60 |
|  |  |  |  | HLA-DRB 4*01:01 | 13.00 |
|  |  |  |  | HLA-DRB 1*15:01 | 17.00 |
|  |  |  |  | HLA-DRB 5*01:01 | 23.00 |
|  |  |  |  | HLA-DRB 3*01:01 | 29.00 |
|  |  |  |  | HLA-DRB 3*02:02 | 29.00 |

### 3.3 Analysis of population coverage

IEDB population coverage tool [23] is used to calculate the population coverage of the predicted epitopes. The result for class I MHC restriction for the China, whole world and Indian population with the selected MHC I alleles is shown in Table 4.

**Table 4.**
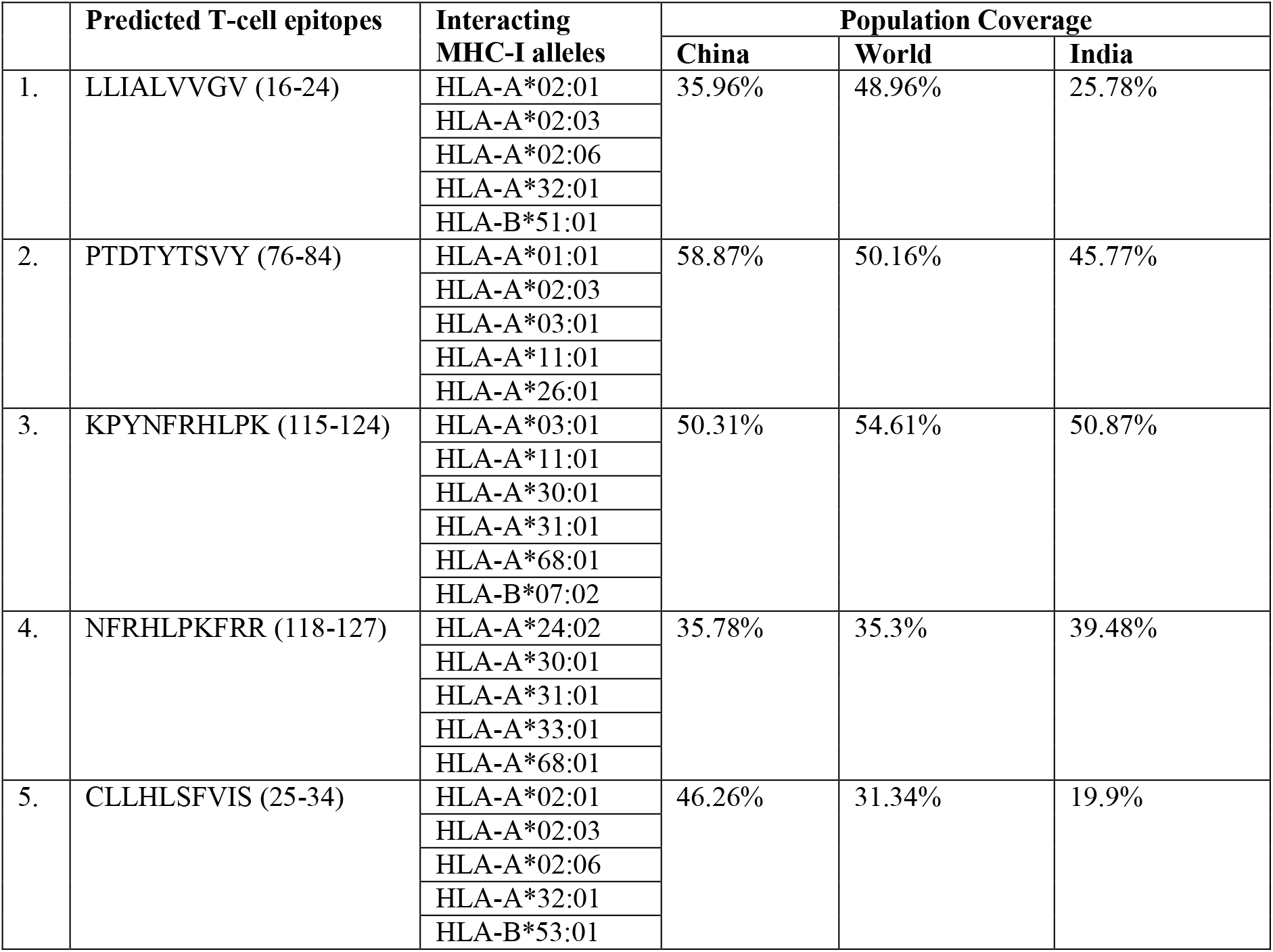
Population coverage for predicted T cell epitopes

### 3.4 Potential B cell epitope prediction

Characteristic features of the B cell epitope contain flexibility, hydrophilicity, surface accessibility and beta-turn prediction. Prediction scores of Chou and Fasman beta turn [16], Emini surface accessibility [12], Kolaskar and Tangaonkar antigenicity [11] and Parker hydrophilicity [13] are plotted (Figure 3). Predicted linear B cell epitopes and the accessibility, hydrophilicity, flexibility and beta-turn prediction score for each residue are summarized in Table 5.

**Figure 3.**
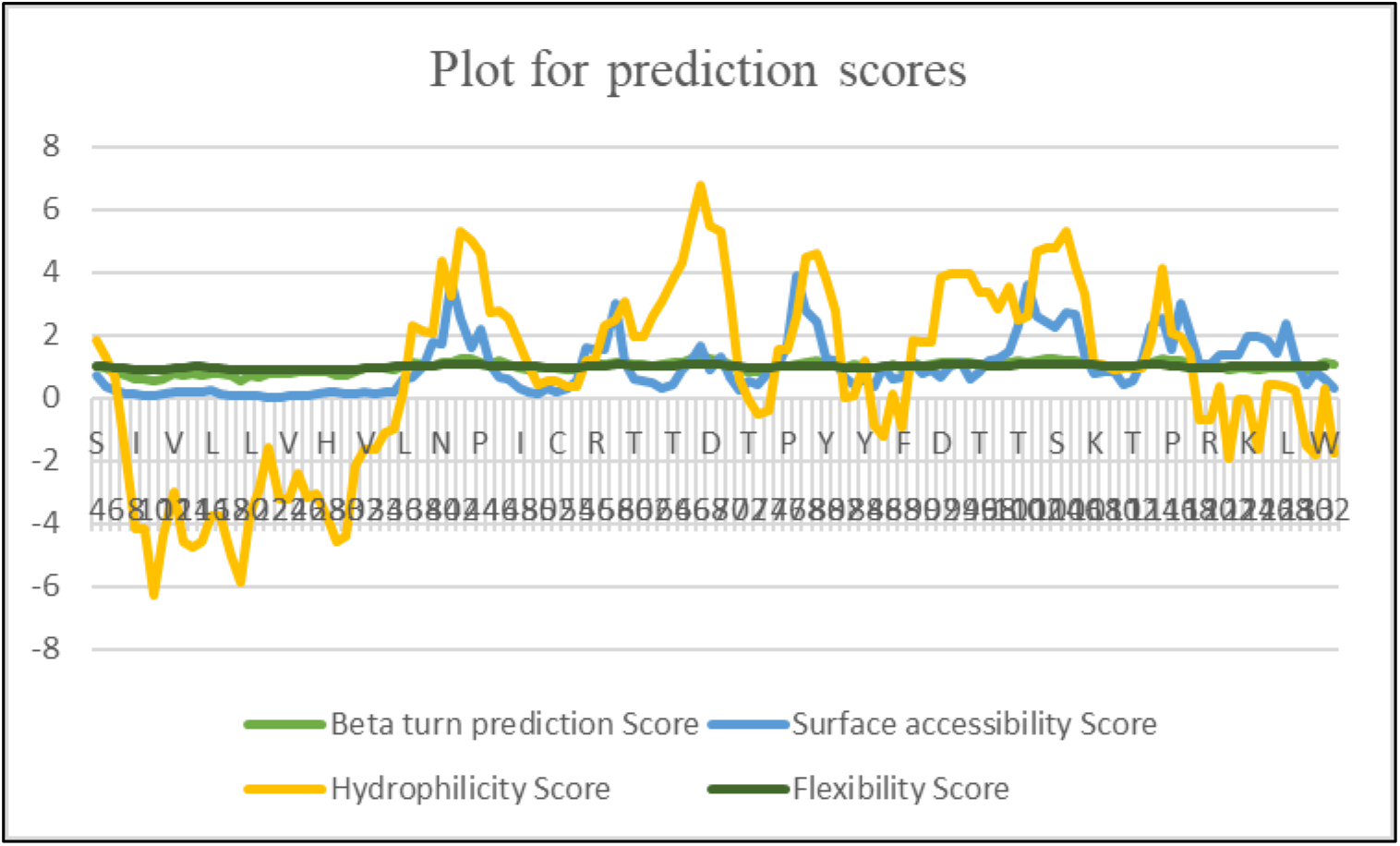
Prediction scores for each amino acid residues of the virus protein

**Table 5.** Surface accessibility, hydrophilicity, flexibility, beta turn and antigenicity prediction score for each residue of B cell epitope.

|  | B cell epitope | Chou and Fashman beta turn score for each residue (threshold=0.993) | Emini surface accessibility score for each residue (threshold=1.000) | Karplus and Schulz flexibility score for each residue (threshold=1.003) | Kolaskar and Tongaonkar antigenicity score for each residue (threshold=1.048) | Parker hydrophilicity score for each residue (threshold=0.740) |
| --- | --- | --- | --- | --- | --- | --- |
| 38 | I | 1.079 | 1.008 | 1.011 | 0.957 | 2.129 |
|  | R | 0.993 | 1.764 | 1.032 | 0.962 | 2.057 |
|  | N | 1.131 | 1.699 | 1.061 | 0.864 | 4.371 |
|  | T | 1.14 | 3.747 | 1.076 | 0.923 | 3.243 |
|  | T | 1.277 | 2.564 | 1.08 | 0.903 | 5.314 |
|  | N | 1.236 | 1.611 | 1.078 | 0.93 | 5.014 |
|  | P | 1.149 | 2.186 | 1.057 | 0.944 | 4.614 |
|  | S | 1.079 | 1.062 | 1.037 | 0.979 | 2.729 |
|  | A | 1.164 | 0.653 | 1.018 | 0.974 | 2.8 |
| 47 | R | 1.079 | 0.61 | 1.001 | 0.993 | 2.543 |
| 56 | R | 1.074 | 1.571 | 1.019 | 1.069 | 1.271 |
|  | T | 1.049 | 1.571 | 1.041 | 0.983 | 2.286 |
|  | P | 1.116 | 3.003 | 1.054 | 0.983 | 2.471 |
|  | P | 1.12 | 1.138 | 1.05 | 1.031 | 3.086 |
|  | T | 1.056 | 0.585 | 1.035 | 1.103 | 1.957 |
|  | S | 1.056 | 0.546 | 1.016 | 1.103 | 1.957 |
|  | V | 1.043 | 0.473 | 1.007 | 1.096 | 2.586 |
|  | V | 1.049 | 0.325 | 1.018 | 1.096 | 3.1 |
|  | T | 1.12 | 0.404 | 1.044 | 1.063 | 3.786 |
|  | S | 1.124 | 0.91 | 1.071 | 1.042 | 4.286 |
|  | G | 1.276 | 1.213 | 1.088 | 0.969 | 5.629 |
|  | D | 1.34 | 1.647 | 1.085 | 0.896 | 6.757 |
|  | D | 1.274 | 0.912 | 1.072 | 0.964 | 5.486 |
|  | G | 1.207 | 1.33 | 1.054 | 0.949 | 5.3 |
|  | R | 1.069 | 0.657 | 1.026 | 1.003 | 3.171 |
| 71 | V | 0.927 | 0.276 | 0.997 | 1.044 | 0.6 |

Prediction scores of Emini surface accessibility, Parker hydrophilicity, Karplus and Schulz
flexibility, Chou and Fashman beta turn for each residue of peptides ^38^IRNTTNPSAR^47^ and ^56^RTPPTSVVTSGDDGRV^71^ reveals that these short stretches of antigenic protein can act as linear B cell epitopes.

### 3.5 Docking study of T cell epitopes

The T cell epitope ^76^ PTDTYTSVYLGKFRG ^90^ (MHC II restricted) is selected on the basis of its interactions with large number of alleles and lowest IC_50_ value with HLA-DRB5*1:01 MHC II allele. Similarly, ^76^ PTDTYTSVY ^84^, restricted with MHC I allelic protein HLA-A*01:01 is designated as most probable T cell epitope which is also present in conserved region of non-structural protein NS4 of coronavirus. Docking studies are performed with these two epitopes with human HLA-DRB5*01:01MHC II molecule and MHC I allelic protein HLA-A*01:01 respectively.

Docking study of PTDTYTSVYLGKFRG epitope with HLA-DRB5*01:01, shows lowest binding energy −786.0 Kcal/mole, shown in Figure 4. Docking structure is stabilized by H bonds, are shown in Table 6.

**Figure 4.**
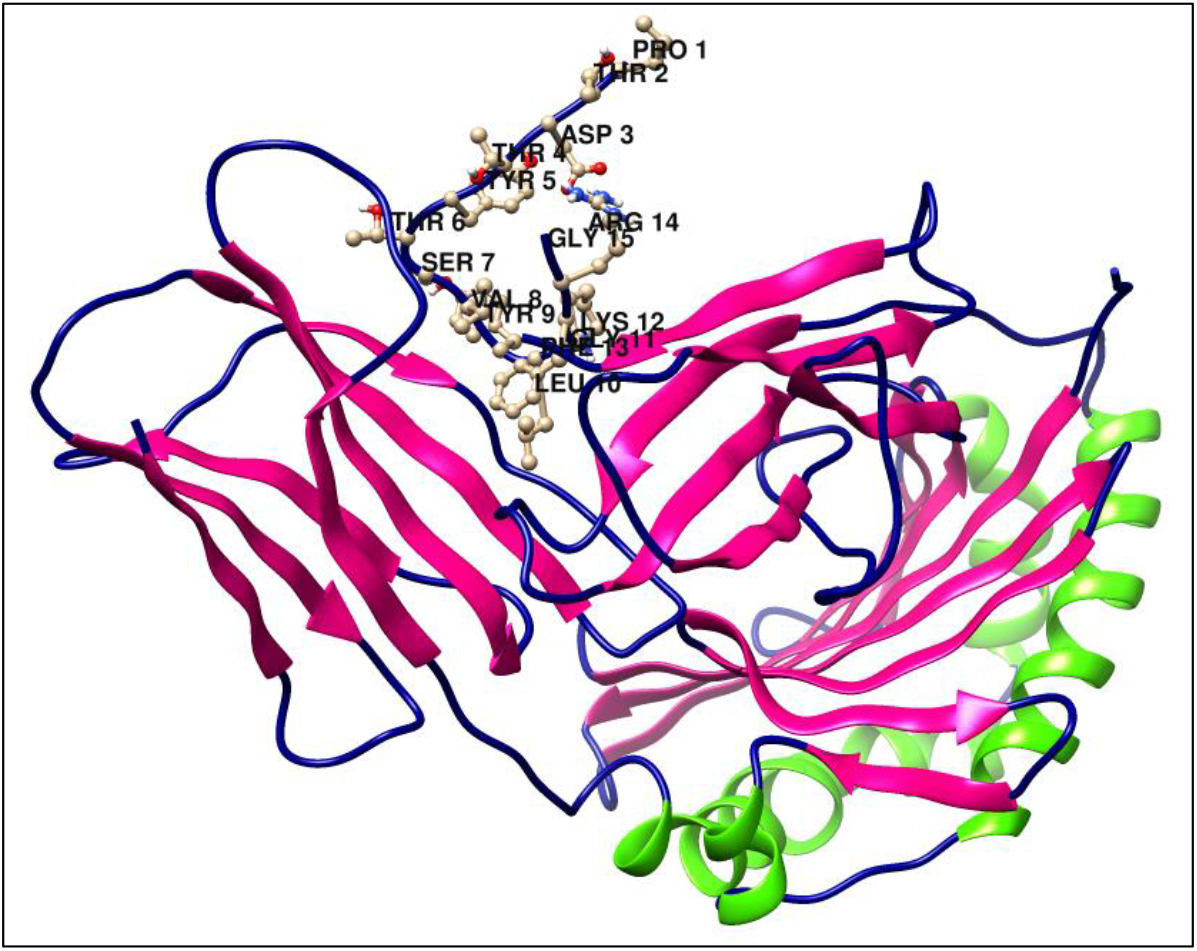
Bound structure of T cell epitope with MHC II

**Table 6.**
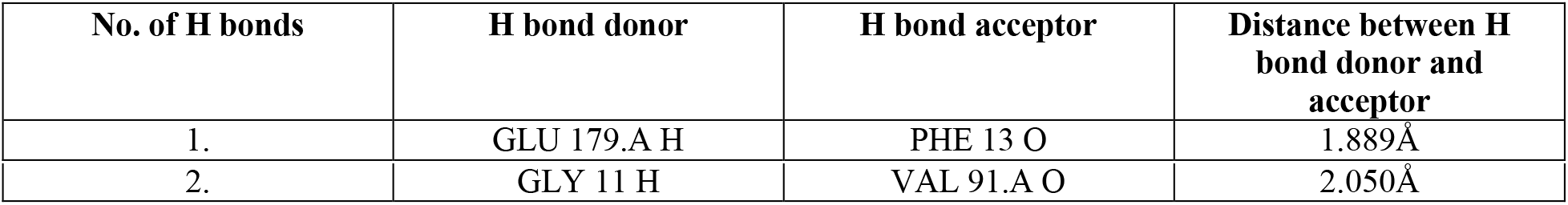
Details of Hydrogen bonding between T cell epitope with MHC II molecule

Interaction between T cell epitope PTDTYTSVY MHC I allelic protein HLA-A*01:01 with binding energy −725.0 Kcal/mole is shown in Figure 5.

**Figure 5.**
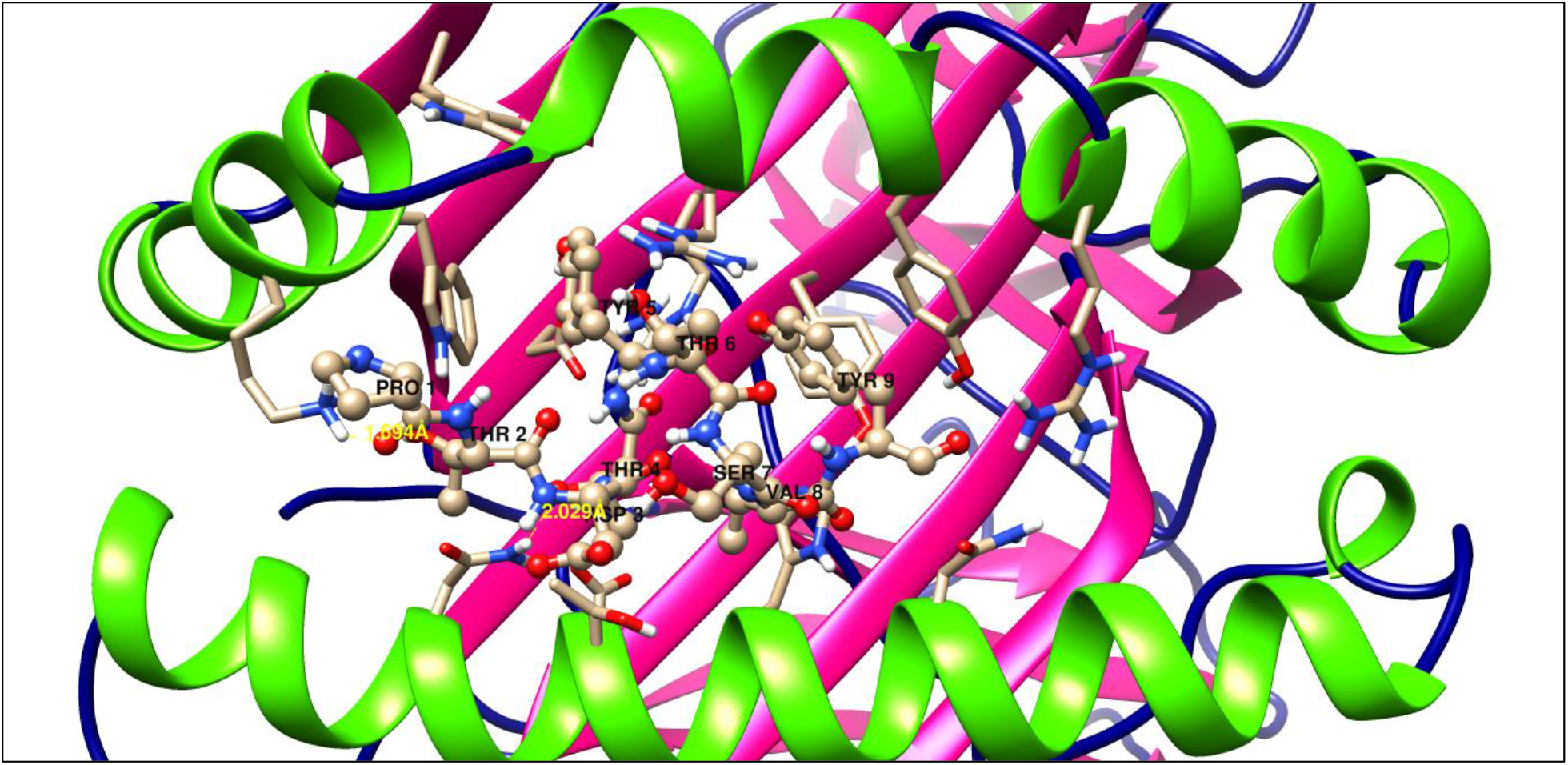
Bound structure of T cell epitope with MHC I molecule

**Table 7.** Details of Hydrogen bonding between T cell epitope with MHC I molecule

| No. of H bonds | H bond donor | H bond acceptor | Distance between H bond donor and acceptor |
| --- | --- | --- | --- |
| 1. | LYS 146.A HZ1 | PRO 1 O | 1.694Å |
| 2. | ASN 77.A HD-22 | THR4 OG1 | 2.029Å |

## 4 Discussions

At present in China and whole world, considering the emergency situation due to corona virus infection, rapid development in vaccine design is the most argent step to prevent pandemics. Because by using vaccine the mortality rate due to coronavirus can be controlled. This technique is applied successfully for smallpox virus, polio virus etc. Though for some other common viruses such as dengue virus, hepatitis C virus, human immunodeficiency virus and coronavirus, vaccine has not been invented till now. Due to lack of definite information about growth, replication and pathogenesis of these viruses. Therefore, computational techniques are used for epitope mapping, which is the preliminary step for vaccine design to prevent coronavirus infection. This study integrates several immunoinformatics and molecular docking methods to recognize potential epitopes of non-structural protein NS4 in coronavirus.

At first, all seventy-nine non-structural protein 4 from different coronaviruses are retrieved from InterPro database and aligned to detect conserved sequences present in them. A phylogenetic investigation reveals a closed evolutionary relationship among these homologous proteins. Non-structural protein 4 is considered as our protein of interest for vaccine design assuming its role in viral replication during coronavirus infection and considering its antigenic nature in human. Not only that, this selected protein has structural similarity with non-structural protein 4 present in rat, murine coronavirus containing various strains. Therefore, the proposed peptide-based vaccine might be effective to prevent coronavirus infection not only in human, but also in rat, murine etc. Here, computational method for vaccine design is totally safe, rapid and cost effective at this emergency situation, caused due to coronavirus infection.

Five potent T cell epitope having potentiality for binding with MHC molecules are predicted. For MHC I and MHC II molecules both 9 mer and 15 mer peptide structures are projected from IEDB recommended prediction method and modeled by peptide modeling algorithm. The percentile rank and IC_50_ values with SMM/ANN method covering all MHC class I supertypes are also analyzed. The five most effective epitopes are presented in Table 2 along with their IC_50_ values.

For MHC I binding prediction scores, the peptide with the lowest percentile rank and IC_50_ value, is selected for their highest affinity for that interacting MHC I allele. The T cell epitope ^76^ PTDTYTSVY ^84^ is considered on the basis of its interaction with large number of alleles and lowest IC_50_ value with HLA-A*01:01 MHC I allele. For this epitope, MHC I processing score with that specific allele comprises proteasome score 1.25, TAP score 1.02, MHC IC_50_ value is 6.7 nm. This means that this specific epitope has high affinity to MHC I molecule during antigen presentation. Moreover that, this epitope when interacting with selective MHC I allele shows the highest population coverage not only for Chinese population, but also has higher population coverage compared to other epitopes, for Indian and whole world population. Henceforth, this epitope is considered as epitope of choice for CD8 ^+^ T cells.

Likewise, presence of wider peptide binding groove in MHC II molecule than that of MHC I, 15 mer epitopes are investigated by smm/nn/sturnilo method along with their IC_50_ values and are listed in Table 3. For MHC II binding prediction method, a 15 mer T cell epitope sequence ^76^ PTDTYTSVYLGKFRG ^90^ of non-structural protein 4, displays a percentile rank 22.0 with IC_50_ value 22.0 at the time of interaction with HLA-DRB5*01:01MHC II allele. This result approves that this peptide can be selected as T cell epitope with MHC II restriction for our protein of interest. Moreover, this T cell epitope sequence is well conserved among the non-structural protein 4 present in different corona viruses.

B cell epitope identification method, the prediction scores for Emini surface accessibility, Parker hydrophilicity, Chou and Fashman beta turn and Karplus and Schulz flexibility for each residue of peptides IRNTTNPSAR, starting from sequence 38 position and ending at 47 position of that viral antigenic protein, predicts that this is the most potent B cell epitope present in it. Additionally, Kolarskar and Tangaonkar antigenicity prediction values, confirm our estimation.

The predicted T cell epitopes are validated by using molecular docking study. These two epitopes are preferably fitted in the epitope binding grooves of the two MHC protein molecules with highly negative binding energies and stabilized by H-bonds.

An important factor in vaccine design is the distribution of selected HLA allelic protein. This distribution varies among the human population according to the population in different geographic regions of the world. Our predicted T cell epitope, ^76^ PTDTYTSVY ^84^, bound with MHC I HLA-A*01:01 allele, which is present among 58.87% and 45.77% of Chinese and Indian populations respectively and 50.16% of world populations. So, it may be concluded that the predicted T cell epitope must be specifically restricted with the predominant MHC molecule, which is present in target population in India and China, as well as in whole world against coronavirus.

## 5 Conclusion

Though in general most peptide-based vaccines are developed considering B cell epitopes, in our present study both B cell and T cell epitopes, present in non-structural protein 4, are considered for vaccine design against coronavirus. These two T cell epitopes can stimulate immunogenic response after administration inside the human body. This immunological reaction can prevent coronavirus infection in human as well as rat and murine, when they come in contact with this virus in future. Since these epitopes are well restricted with MHC molecules and at the same time, they are almost conserved among other homologous proteins. These proteins include non-structural protein 4, obtained from novel-coronavirus infected patient in China. Thus, these epitopes can be proceeded for further experimental verification during vaccine designing against coronavirus infection.

